# Sequential rescue and repair of stalled and damaged ribosome by bacterial PrfH and RtcB

**DOI:** 10.1101/2021.09.22.461353

**Authors:** Yannan Tian, Fuxing Zeng, Adrika Raybarman, Amy Carruthers, Qingrong Li, Shirin Fatma, Raven H. Huang

**Affiliations:** Department of Biochemistry, University of Illinois at Urbana-Champaign, Urbana, IL 61801, USA; Department of Biology, Southern University of Science and Technology, Shenzhen, P.R. China

**Author notes:** These authors contribute equally: Yannan Tian, Fuxing Zeng, Adrika Raybarman.

## Abstract

In bacteria, rescue of stalled ribosomes due to 3’-truncated mRNAs is carried out by the ubiquitous trans-translation system as well as alternative ribosome-rescue factors such as ArfA and ArfB. It is unclear, however, how the stalled ribosomes caused by ribosomal damages are rescued. Here, we report that a bacterial system composed of PrfH and RtcB not only rescues a stalled ribosome resulting from a specific damage in the decoding center but also repairs the damage afterwards. Peptide release assays reveal that PrfH is only active with the damaged ribosome, but not with the intact one. A 2.55-angstrom cryo-EM structure of PrfH in complex with the damaged 70S ribosome provides molecular insight into specific recognition of the damage site by PrfH. RNA repair assays demonstrate that PrfH-coupled RtcB efficiently repairs the damaged 30S ribosomal subunit, but not the damaged tRNAs. Thus, our studies have uncovered a biological operation by a pair of bacterial enzymes, aiming to reverse the potentially lethal damage inflicted by an invading ribotoxin for cell survival.

## Introduction

Conflicts between organisms are fundamental biological phenomena. In an environment of limited nutrition, some bacteria produce small-molecule or protein toxins to kill their neighbors to eliminate competition. Because the protein translation machinery is essential and universally conserved in all organisms, it is the main target of both small-molecule and protein toxins. The majority of protein toxins targeting protein translation are ribotoxins, inflicting damage on various RNAs required for protein translation. The major sites targeted by ribotoxins include the sarcin-ricin loop (SRL) in the large ribosomal subunit^1,2^, the decoding center in the small ribosomal subunit^3–6^, and the anticodon loops of tRNAs^7–11^. Of particular relevance to this work is the damage done in the decoding center. To date, two different families of ribotoxins, the C-terminal toxin domain of colicin E3 (ColE3-CT) and the C-terminal toxin domain of CdiA effector from *Enterobacter cloacae* (CdiA-CT^ECL^) cleave 16S rRNA at the same site (between nucleotides A1493 and G1494), demonstrating convergent evolution of different lineages of ribotoxins targeting one of the most conserved sites in the ribosome.

To survive, the bacteria targeted by toxins employ a variety of defensive mechanisms of neutralizing the toxins imposed on them. The majority appear to employ a head-on approach to directly confront the invading toxins. Here we propose that, if the damage inflicted by an invading toxin is reversible, an alternative approach of indirect confrontation could also be employed. The clash between a ribotoxin and an RNA repair system appears to be such an example, as the conflict is carried out via an RNA intermediate. We have previously reported two bacterial RNA repair systems that are able to repair ribotoxin-cleaved tRNAs *in vitro*^12,13^. Here we explore possible RNA repair by bacterial RtcB^14–16^. In archaeal and most eukaryotic organisms, RtcB is the ligase component of the tRNA splicing complex^17,18^. However, most RtcB are found in bacteria and bacterial tRNAs have no standard introns. Because tRNAs are the substrates of archaeal and eukaryotic RtcB, it is assumed that bacterial RtcB is involved in repairing damaged tRNAs. Here we provide experimental evidence to demonstrate that, in addition to the damaged tRNAs, bacterial RtcB also carries out repair of damaged ribosomes.

## Results

### The 70S ribosome damaged in the decoding center is the substrate of PrfH

To provide insight into biological functions of RtcB, we first performed bioinformatic analysis of Pfam PF01139, the RtcB protein family. Our analysis revealed that the vast majority of RtcB are indeed found in bacteria (Supplementary Fig. 1a). We also constructed a Sequence Similarity Network (SSN) of PF01139 (Supplementary Fig. 1b). SSN revealed that a subset of RtcB, denoted RtcB2 here, forms a separate cluster (Supplementary Fig. 1b, cluster 2), suggesting that RtcB2 might be functionally distinct from the majority of RtcB from cluster 1. Furthermore, the overwhelming majority of bacterial RtcB2 are encoded in a two-component operon that also encodes PrfH^19^ (Supplementary Fig. 2a). Both RtcB2 and PrfH have not been experimentally characterized before.

To elucidate biological function of bacterial PrfH, we cloned, overexpressed, and purified recombinant PrfH from *E. coli* (*Ec*PrfH). When the intact *E. coli* 70S ribosome and a 25-nucleotide mRNA (Fig. 1a, mRNA-25) were employed as the substrates, no enzymatic activity of *Ec*PrfH was detected (Fig. 1b, red curve). We reasoned that, since *Ec*PrfH and *E. coli* RtcB2 are encoded in the same operon (Supplementary Fig. 2), a damaged ribosome might be required for the activity of *Ec*PrfH. We therefore prepared and purified *E. coli* 70S ribosome having 16S rRNA cleaved by CdiA-CT^ECL^ to test our hypothesis. The peptide release assays using the damaged 70S ribosome indeed showed robust activity of *Ec*PrfH (Fig. 1b, green curve), indicating that 70S ribosome with a specific damage in the decoding center is likely the biological substrate of PrfH.

**Fig. 1.**
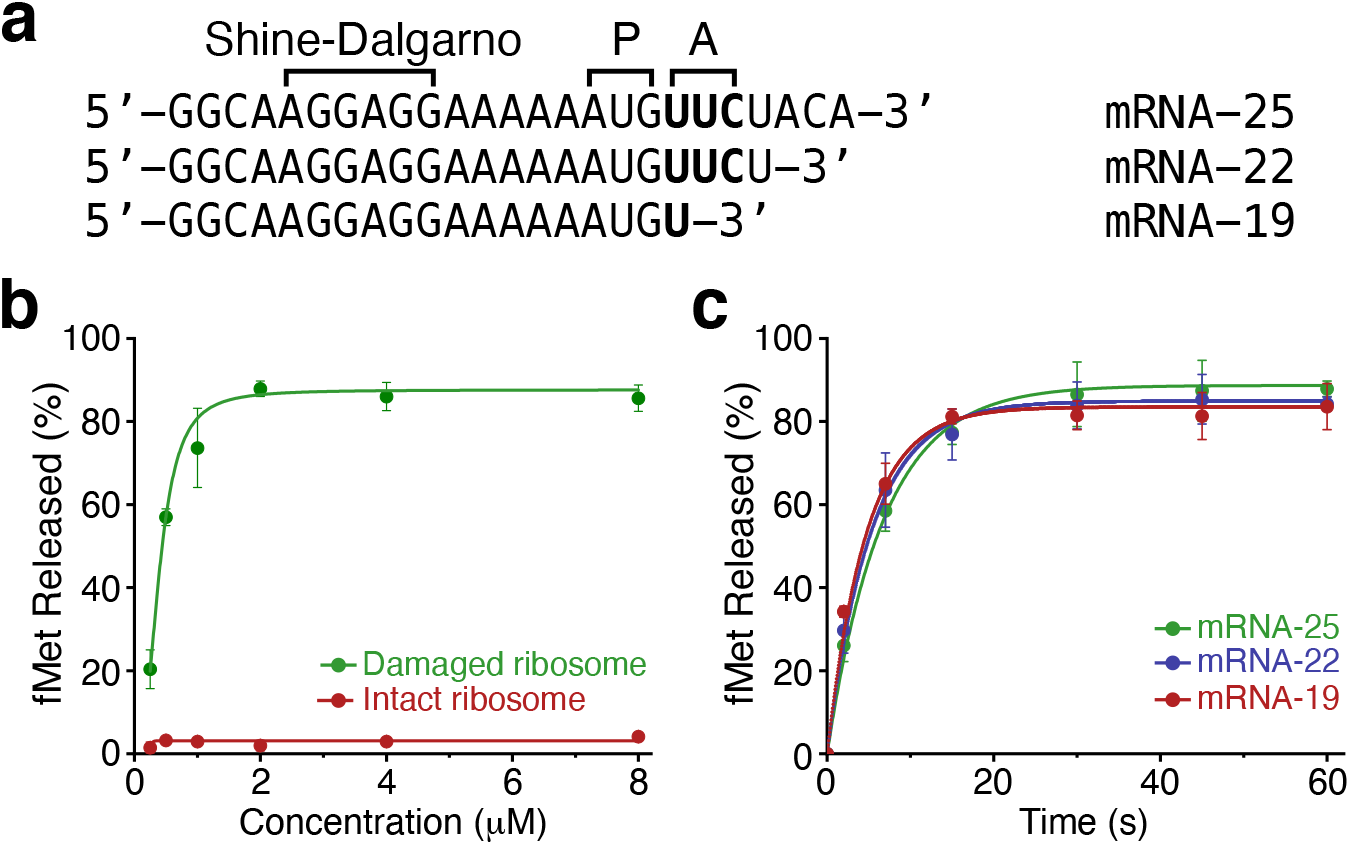
Substrate specificity of peptide release by *Ec*PrfH. **a** Sequences of three mRNAs with different length employed for the assays. P, P site of ribosome; A, A site of ribosome. **b** Concentration-dependent activities of PrfH using the damaged 70S ribosome (green) and the intact 70S ribosome (red) as substrates. mRNA-25 was employed for the assays. **c** Time course of the activities of PrfH with three different mRNAs. The concentration of PrfH employed for the assays is 2 μM.

Bacterial RF1 and RF2 require the presence of mRNA with stop codons in the A site. To assess whether the activity of PrfH also requires mRNA in the A site, we prepared two additional mRNAs with systematic truncation at their 3’-ends that encompass the A site (Fig. 1a). PrfH is approximately equally active with all three mRNAs (Fig. 1c), indicating that, unlike RF1 and RF2, the activity of PrfH does not depend on the presence of mRNA in the A site. We carried out additional peptide release assays, allowing us to obtain kinetic parameters of the PrfH-catalyzed reaction (Supplementary Fig. 3).

### Cryo-EM structure of the damaged *E. coli* 70S•PrfH•tRNA•mRNA complex

To provide molecular insight into the recognition of the damaged 70S ribosome by PrfH, we determined the cryo-EM structure of *Ec*PrfH in complex with the damaged *E. coli* 70S ribosome, tRNA^fMet^, and mRNA-25 at 2.55 Å resolution (Fig. 2a, Supplementary Figs. 4, 6, Supplementary Table 1). The structure revealed the presence of mRNA (Fig. 2a, blue) as well as tRNA^fMet^ occupying both the P and E sites of the ribosome (Fig. 2a, green and marine). Importantly, *Ec*PrfH is found to occupy the A site approximately where RF1 or RF2 should locate during normal translation termination^20,21^ (Fig. 2a, colored red). We also carried out a similar structural study with the intact *E. coli* 70S ribosome, and no *Ec*PrfH could be found in the A site (Supplementary Fig. 5). This is consistent with the result of peptide release assays that the intact 70S ribosome is not a substrate of PrfH (Fig. 1b).

**Fig. 2.**
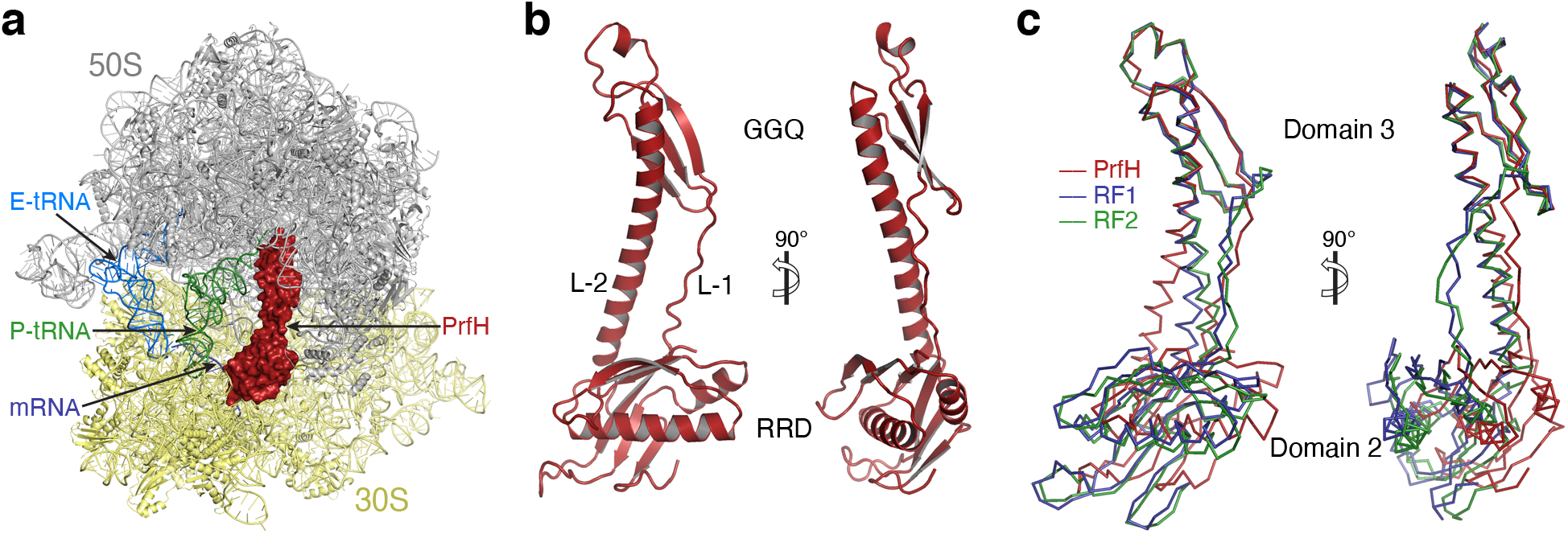
Structure of *Ec*PrfH in complex with the damaged *E. coli* 70S ribosome, P- and E-site tRNAs, and mRNA-25. **a** Overview of the structure of the complex. Two subunits of the ribosome are colored gray and yellow, respectively. P-site tRNA is colored green, E-site tRNA is colored marine, and mRNA is colored blue. PrfH occupies the A site of the damaged ribosome and is depicted in surface and colored red. **b** Cartoon representation of the PrfH structure. RRD, rRNA-recognition domain; GGQ, peptide-hydrolase domain that contains the strictly conserved GGQ motif; L-1 and L-2, linker-1 and linker-2. **c** Ribbon representation of the structures of PrfH (red), RF1 (blue), and RF2 (green), aligned based on the superimpositions of the structures of 70S ribosomes associated with these factors.

The folding of *Ec*PrfH is similar to the structures of domain 2, domain 3, and part of domain 4 of RF1 and RF2 (Fig. 2b, c, Supplementary Fig. 7). This is further confirmed by the Dali structural search^22^ with *Ec*PrfH, as the top scores are all structures of RF1 and RF2. To facilitate structural analysis of *Ec*PrfH, we divided the structure of *Ec*PrfH into three portions: the top and bottom domains and the linkers connecting these two domains (Fig. 2b, Supplementary Fig. 8). The top domain is composed of residues 108-161, and this domain is highly homologous to domain 3 of RF1 and RF2 (Supplementary Fig. 7a). This domain is therefore named GGQ domain (Fig. 2b, top). The bottom is composed of residues 2-98 and residues180-204. As described in the next section, this domain is responsible for the recognition of ribosomal RNAs, and it is therefore named rRNA-recognition domain (RRD) (Fig. 2b, bottom). These two domains are connected by two linkers, a loop composed of residues 99-107 (Linker-1), and part of a long helix composed of residues 162-179 (Linker-2) (Supplementary Fig. 8).

The structure of PrfH is highly homologous to the ones of RF1 and RF2 when individual domains are compared (Supplementary Fig. 7a, b), but less so when the entire structures are compared (Supplementary Fig. 7c). The cause of this disparity is the changes of relative position and orientation of the two domains in PrfH compared to their counterparts in RF1 and RF2. This is clearly demonstrated when PrfH, RF1, and RF2 are aligned based on the superpositions of the 70S ribosome structures that are associated with these factors (Fig. 2c). While the GGQ domain of PrfH and domain 3 of RF1 and RF2 are in the same position, the rRNA-recognition domain of PrfH shifts ∼8 Å vertically, ∼16 Å horizontally, and rotates ∼25º vertically, relative to domain 2 of RF1 and RF2 (Fig. 2c). As a result, the rRNA-recognition domain of PrfH no longer makes contacts with mRNA (Supplementary Fig. 9). Instead, it interacts extensively with rRNAs as well as S12 protein, made possible with the breakage of the phosphate backbone between A1493 and G1494 by CdiA-CT^ECL^.

### Molecular recognition of the damaged 70S ribosome by PrfH

In the decoding center of bacterial ribosome, several universally conserved residues are responsible for monitoring codon-anticodon interactions^23^. They are A1492 and A1493 from helix 44 (h44), C518 and G530 from helix 18 (h18), C1054 from helix 34 (h34), and S47 from ribosomal protein S12. Except for A1492, all these residues make direct contacts with PrfH (Fig. 3a). In addition, residue C1914 from helix 69 (H69) of 23S rRNA also interacts with PrfH (Fig. 3a, colored green).

**Fig. 3.**
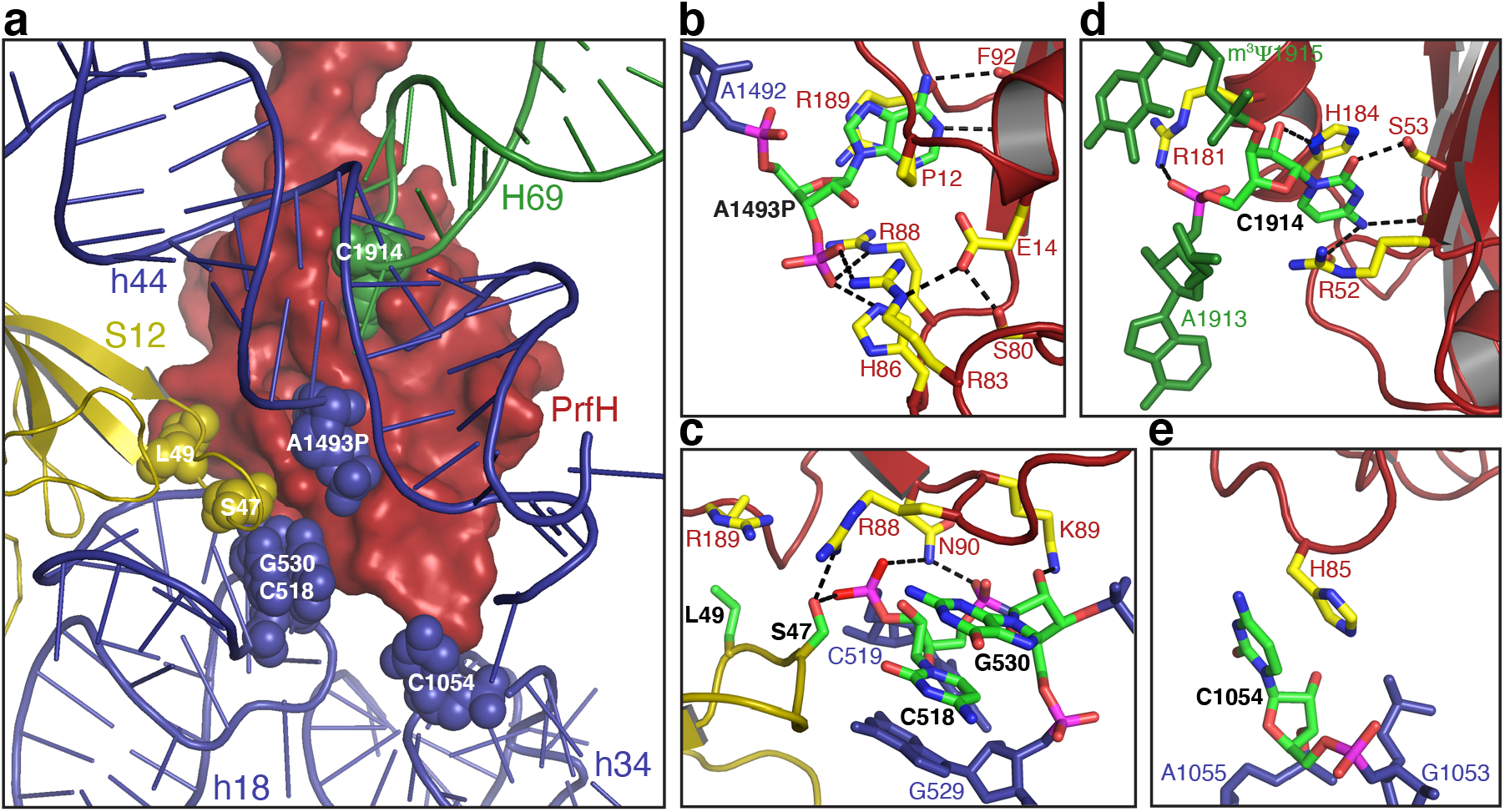
Molecular recognition of the damaged 70S ribosome by PrfH. **a** Overview of the nucleotides and the amino acids in the damaged 70S ribosome making direct contacts to PrfH. PrfH is in surface, and the ribosome is in cartoon. The residues that make direct contacts to PrfH are highlighted in spheres. **b** Recognition of the 3’-terminal A1493P of the cleaved 16S rRNA by PrfH. Atoms in A1493P and in the residues from PrfH are colored individually, with carbon in green (A1493P) and yellow (PrfH), oxygen in red, nitrogen in blue, and phosphate in magenta. **c** Specific interactions between PrfH and residues from h18 of 16S RNA and S12 protein. **d** Recognition of C1914 of 23S rRNA by PrfH. **e** Observed single interaction between H85 of PrfH and C1054 from h34 of 16S rRNA.

The center of the rRNA-recognition domain is a pocket that specifically recognizes the 3’-terminal nucleotide of the cleaved 16S rRNA: 3’-phosphorylated A1493 (denoted A1493P here). The A1493P-recognition pocket is mainly surrounded by two walls (Supplementary Fig. 10d). The base of A1493P is sandwiched between the side chain of R189 and residues G10-P11, and the ribose stacks on top of the side chain of R88 (Fig. 3b). The base of A1493P is recognized via two hydrogen bonds with the main chain of F92. Perhaps the most extensive interactions occur at the 3’-phosphate group that originally belongs to G1494 prior cleavage. The side chains of three residues, R83, H86, and R88, form hydrogen bonds with the phosphate group (Fig. 3b). The side chain of R83 is stabilized by a hydrogen-bonding network that also involves E14 and S80.

On the other side of the first wall is a highly conserved surface patch that is responsible for interactions with residues from h18 and S12 protein (Fig. 3c, Supplementary Fig. 10d). Specifically, the side chain of N90 hydrogen bonds with the phosphate backbones of both C518 and C519, and the side chain of K89 hydrogen bonds with the 2’-OH group of G530 (Fig. 3c). The side chain of R88 hydrogen bonds with the side chain of S47 of S12 protein, which in turn hydrogen bonds with the phosphate of C519. In addition, the side chain of R189 stacks on the side chain of L49 of S12 protein.

On the other side of the second wall is a second pocket for specific recognition of C1914 from H69 (Fig. 3d). Specifically, the base of C1914 is sandwiched between the side chains of H184 and R52. H184 also hydrogen bonds with the 2’-OH group of C1914. The base of C1914 is recognized by three hydrogen bonds: one between O2 and the side chain of S53, and two between N4 and the carbonyl groups of A9 and R52 (Fig. 3d). In addition, the side chain of R181 hydrogen bonds with the phosphate group of C1914, and it also stacks on m^3^Ψ 1915. PrfH also makes a single contact with C1054 from h34 via stacking of the side chain of H85 on the base of C1054 (Fig. 3e).

PrfH binding results in significant conformational changes of rRNAs of the damaged 70S ribosome (Supplementary Fig. 11). Notably, because A1493P is flipped out, A1913 from H69 occupies the space vacated by A1493P, stacking on top of A1492 (Supplementary Fig. 11a). The conformations of rRNAs of the PrfH structure are significantly different from the ones when tRNA^24^ or RF2^21^ occupies the A site (Supplementary Fig. 11b), but they share some similarities with the structures when ArfA•RF2 or ArfB occupies the A site^25–32^ (Supplementary Fig. 11c).

### *In vivo* RNA substrates of *Ec*RtcB1 and *Ec*RtcB2 are mutually exclusive

In addition to the well characterized *E. coli* RtcB (denoted *Ec*RtcB1 here), the founding member of RtcB family, all *E. coli* strains encode a second RtcB (denoted *Ec*RtcB2 here). These two RtcB only share 28% sequence identity, implying their distinct biological functions. While *Ec*RtcB1 is encoded in a two-component operon that also encodes RtcA^33^, *Ec*RtcB2 is encoded in a two-component operon that also encodes *Ec*PrfH (Supplementary Fig. 2). In *E coli* K-12 strain and its derivatives, RtcB2 and PrfH are presumably inactive due to a significant genomic deletion that encompasses both genes (Supplementary Fig. 2b). In addition to these two RtcB, approximately 60 *E. coli* strains encode a third RtcB (*Ec*RtcB3) that is distinct from both *Ec*RtcB1 and *Ec*RtcB2.

To provide insight into substrate specificity of *Ec*RtcB2, we prepared three ribotoxin-cleaved *E. coli* RNA substrates for qualitative RNA repair assays (Supplementary Fig. 12). They are tRNA^Asp^ cleaved by the C-terminal toxin domain of colicin E5 (ColE5-CT) between nucleotides 34 and 35, tRNA^Arg^ cleaved by the C-terminal toxin domain of colicin D (ColD-CT) between nucleotides 38 and 39, and 30S subunit having 16S rRNA cleaved by CdiA-CT^ECL^. In addition to recombinant *Ec*RtcB2 potentially repairing these three RNA substrates, we also carried out identical sets of experiments using recombinant *Ec*RtcB1 for comparison. It is surprising that, despite extensive biochemical characterization of *Ec*RtcB1, no cleaved RNA substrates isolated from *E. coli* have been tested *in vitro* for potential RNA repair by this enzyme.

*Ec*RtcB1 efficiently repairs damaged tRNA^Asp^ (Fig. 4a and Fig. 4g, blue curve). Although the final yield of repair with the damaged tRNA^Arg^ is not as high as the one with the damaged tRNA^Asp^, the initial rates of repair are comparable with both damaged tRNAs (Fig. 4g, blue and green curves at the reaction time points of 2 and 5 minutes). Therefore, the modest repair yield of the damaged tRNA^Arg^ might result from the poor quality of the RNA substrate. This is consistent with the observed degradation of tRNA^Arg^ substrate over time (Fig. 4c, d). Based on the quantitation of full-length 16S rRNA and 3’-cleaved fragment (Fig. 4e), less than 3% of the damaged 30S subunit has been repaired by *Ec*RtcB1 at its maximum (Fig. 4g, red curve with the reaction time point of 5 minutes). This could be caused by the uncertainty of quantitation. Alternatively, the number might represent *Ec*RtcB1 repairing some very minor non-specific ribosomal damages outside the decoding center where *Ec*RtcB1 might have access. *Ec*RtcB1 has previously been shown to reduce non-specific rRNA fragmentation resulting from cell stress^34^. We argue that, if it were from the repaired product of 30S subunit at the site cleaved by CdiA-CT^ECL^, one would have expected the increase of repair yield over reaction time. This is clearly not what happened (Fig. 4g, red curve). Therefore, we conclude that *Ec*RtcB1 does not repair 30S subunit with the specific damage inflicted by CdiA-CT^ECL^.

**Fig. 4.**
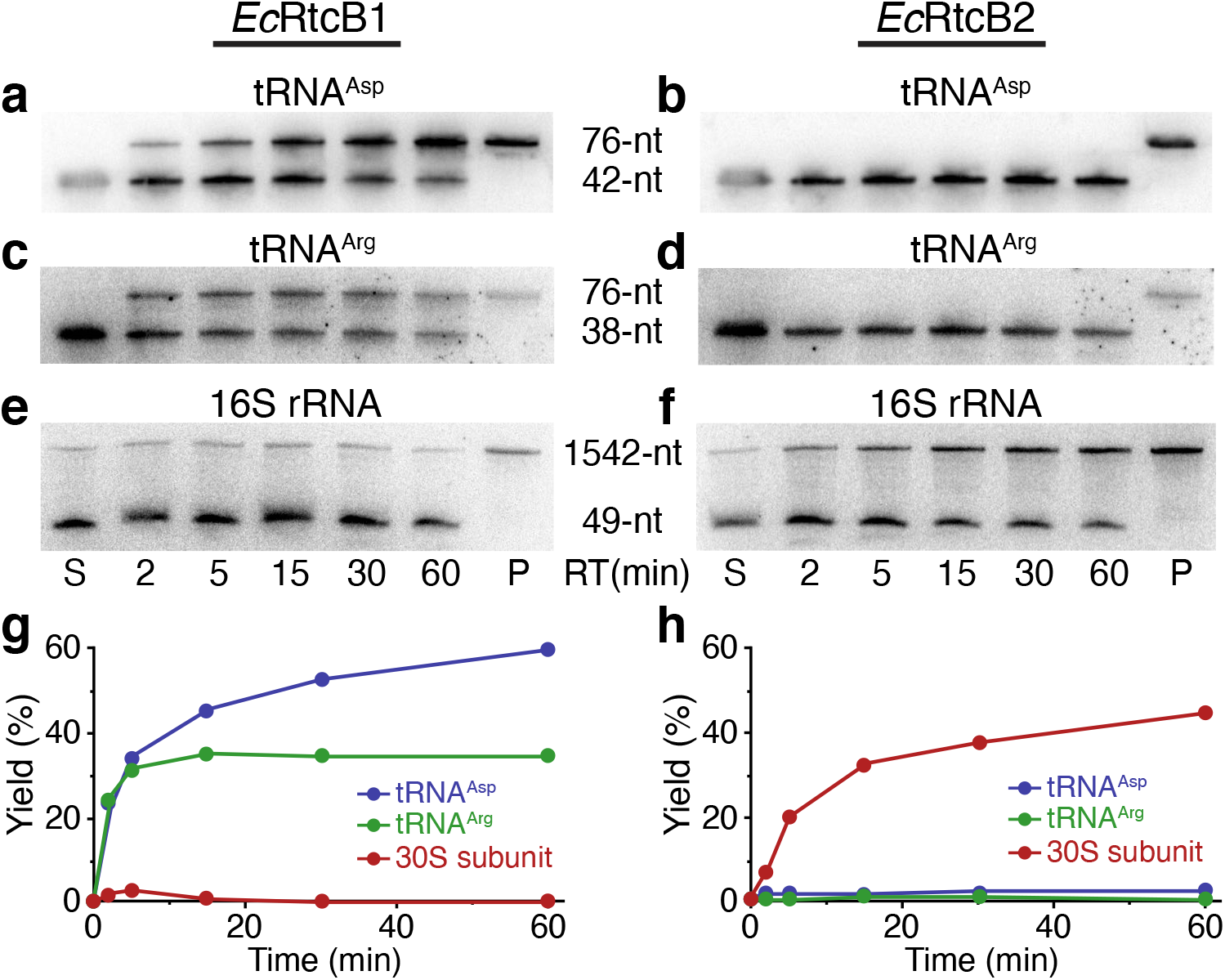
Substrate specificity of RNA repair by *Ec*RtcB1 and *Ec*RtcB2. **a-f** Northern blotting of denaturing polyacrylamide gels analyzing RNA repair of three different RNA substrates by *Ec*RtcB1 and *Ec*RtcB2. S, substrates; P, total RNAs isolated from *E. coli* cells without the exposure to ribotoxins; RT, reaction time. **g** Quantitation of the RNA repair reactions carried out by *Ec*RtcB1. **h** Quantitation of the RNA repair reactions carried out by *Ec*RtcB2.

The results of RNA repair by *Ec*RtcB2 are just the opposite. *Ec*RtcB2 efficiently repairs damaged 30S subunit (Fig. 4f and Fig. 4h, red curve), but does not repair damaged tRNAs (Fig. 4b, d). Again, less than 2% repair yield with the damaged tRNA^Asp^ might be caused with the uncertainty of quantitation. Indeed, we did not observe any bands corresponding to the repaired products when we re-analyzed only the portion of the gel corresponding to the hypothetical repair products (data not shown). Based on these analyses, we conclude that *Ec*RtcB2 does not repair damaged tRNAs. The contrast of pair-wise comparisons of the gels shown in Fig. 4 is striking, reflecting completely different substrate specificity of two RtcB present in *E. coli*. Therefore, if the results of our *in vitro* RNA repair assays stand *in vivo*, the biological roles of *Ec*RtcB1 and *Ec*RtcB2 are different and mutually exclusive. *Ec*RtcB1 is mainly responsible for repairing damaged tRNAs, whereas *Ec*RtcB2, with the help from *Ec*PrfH, specifically repairs the ribotoxin-induced cleavage of 16S RNA between A1493 and G1494.

## Discussion

Based on the biochemical and structural data presented here, we propose the following biological events occurring in bacteria that have been invaded by a ribotoxin such as ColE3-CT or CdiA-CT^ECL^. Ribosomal damage in the decoding center results in the stalled 70S ribosome (Fig. 5, first panel). Unlike the stalled ribosomes due to 3’-truncated mRNA, the stalled ribosome caused by ribosomal damage should still have mRNA occupying the A site and beyond. Furthermore, the population of the stalled ribosome should be significantly higher than the one due to 3’-truncated mRNAs, which was estimated to be 2-4%^35^. RNA repair cannot occur with the damaged 70S ribosome as RtcB2 is not able to access the damage site. PrfH enters the empty A site (Fig.5, Step 1). Specific interactions of PrfH with rRNAs and S12 protein of the damaged ribosome orients its GGQ domain to catalyze hydrolysis of the nascent peptide attached to P-site tRNA (Fig. 5, Step 2). The removal of the nascent peptide allows 70S ribosome to split into 50S and 30S subunits (Fig. 5, Step 3). It is unclear whether PrfH alone is sufficient for the split of the 70S ribosome, or additional cellular factors are required. RtcB2 accesses the damage site of the dissociated 30S subunit, performing RNA repair using GTP as the cofactor (Fig. 5, Step 4). The repaired 30S subunit re-enters the ribosome pool, assembling into a new 70S ribosome to start a new round of protein translation (Fig. 5, Step 5).

**Fig. 5.**
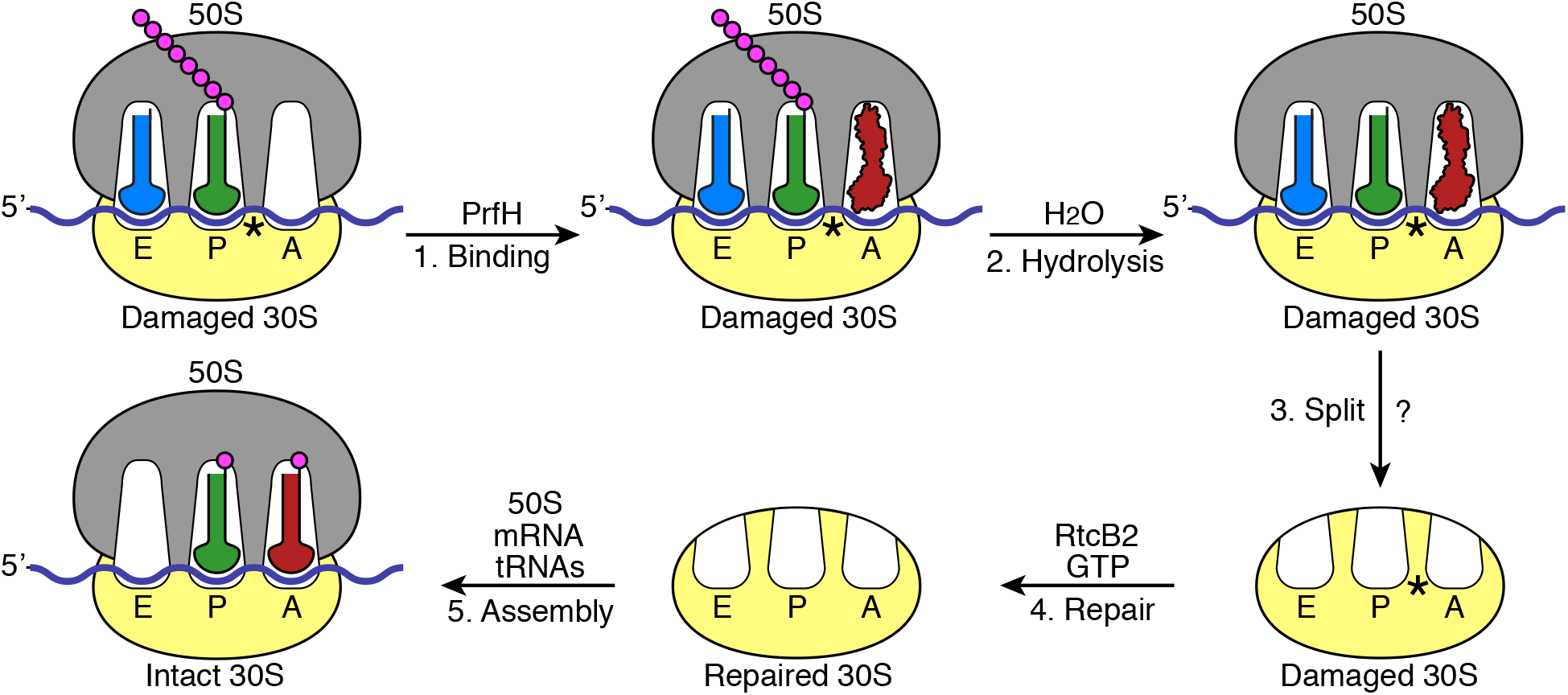
Proposed biological events occurring in a bacterial cell invaded by a ribotoxin such as ColE3-CT or CdiA-CT^ECL^. The protein translation machinery is schematically depicted, including 50S ribosomal subunit (gray), 30S ribosomal subunit (yellow), mRNA (blue), A-, P- and E-site tRNAs (red, green, and marine, respectively), amino acids (magenta), and PrfH (red). The ribosomal damage caused by a ribotoxin is marked with an asterisk.

Because of extensive and precise interactions between PrfH and the damaged 70S ribosome revealed by the cryo-EM structure (Fig. 3), it is difficult to envision that PrfH is capable of rescuing stalled 70S ribosomes other than the one described here. Therefore, PrfH appears to have evolved to deal with only one problem: the biological consequence resulting from specific cleavage of 16S rRNA between A1493 and G1494. Recently, Aravind and coworkers performed a comprehensive bioinformatic analysis of the origin and evolution of release factors^36^. The authors assigned PrfH a separate branch on the phylogenetic tree in parallel with the branch that belongs to RF1 and RF2. Although our structural comparison revealed that the rRNA-recognition domain of PrfH is homologous to domain 2 of both RF1 and RF2, it fits better with RF2 based on the parameters of rmsd and sequence identities (Supplementary Fig. 7b). In addition, the SPF motif of RF2, critical for the stop codon recognition, is semi-conserved in PrfH, with strict conservation of the first residue (Supplementary Fig. 8, boxed in red). While Serine of the SPF motif in RF2 makes a direct contact with the second base of the stop codon^21^, the equivalent residue in PrfH participates in recognition of the terminal A1493P (Fig. 3b, S80). In fact, the loop where this motif resides makes the biggest contribution to recognizing the damaged 70S ribosome, supplying six out of eleven residues that make direct contacts with the damaged 70S ribosome (Fig. 3). Therefore, it is possible that bacterial PrfH might have evolved from RF2, with the loss of domain 1, the loss of part of domain 4, and several mutations that convert the codon-recognition domain into the rRNA-recognition domain.

Colicin E3 is the first ribotoxin to be characterized 50 years ago^3,4^, disabling 70S ribosome for protein translation via specific cleavage of 16S rRNA between A1493 and G1494. Over the last half a century, it was unclear whether such a lethal ribosomal damage could be reversed to allow cell to survive. Here we demonstrate that it can be done through the combined effort of PrfH and RtcB2. In addition, our RNA repair assays reveal that RtcB2 is strictly specific for repairing 30S subunit with a specific cleavage of 16S rRNA between A1493 and G1494. Therefore, although we cannot rule out the possibility that RtcB2 might have enzymatic activity on other RNA substrates *in vitro*, its biological role appears to repair the only ribosomal damage occurring in the decoding center. This argument is certainly consistent with the biological role of PrfH discussed above. Conversely, the fact that many bacteria require the presence of an operon encoding RtcB2 and PrfH underscores the severity of the threat posed by ribosome-specific ribotoxins such as ColE3-CT and CdiA-CT^ECL^ to bacteria.

In addition to bacterial RtcB2, 98 RtcB2 are also present in eukaryotic organisms based on our bioinformatic analysis (Supplementary Fig. 1), and most of those organisms are Fungi and Plants. Since Fungi and Plants employ Trl1 for tRNA splicing^37,38^, we predict that eukaryotic RtcB2 carry out repair of 40S subunit with the damage similar to the one observed here in bacteria. This raises two interesting questions regarding how eukaryotic ribosome is damaged and how the damaged 80S ribosome is rescued, as ribotoxins equivalent to ColE3-CT and CdiA-CT^ECL^ have not been discovered in eukaryotes and PrfH is only found in bacteria. Our analysis also indicates that bacterial RtcB are more diverse than we anticipated. For example, SSN of the entire RtcB family classifies RtcB into eight clusters (Supplementary Fig. 1b). The current study, together with previous studies on archaeal and human RtcB, only allows us to tentatively assign RNA substrates of RtcB from clusters 1a, 1b, and 2. Therefore, it would be interesting to investigate possible biological substrates of RtcB from other five clusters, which might yield new surprises.

## Methods

### Bioinformatic analysis of Pfam PF01139, the RtcB protein family

Bioinformatic analyses were performed on the database of UniProt 2020_04 and InterPro 81. Calculations were carried out at the EFI website (https://efi.igb.illinois.edu/)^39^. PF01139 (Pfam of RtcB) was submitted for the initial calculation for the Sequence Similarity Network (SSN) of RtcB. After the initial calculation was complete, SSN was finalized with the setting of Alignment Score Threshold = 200 and Minimum Sequence Length = 350. The SSN file with 100% ID, e.g., 100% rep node, was displayed with Cytoscape^40^ to produce the initial SSN. The nodes corresponding to Bacteria, Archaea, Eukaryota, and Viruses were selected separately, and SSN corresponding to these four classifications were generated. These four additional SSN allowed us to obtain the numbers of unique sequences of all RtcB for Bacteria, Archaea, Eukaryota, and Viruses as shown in Supplementary Fig. 1a. To obtain the corresponding numbers for RtcB2, similar analysis was carried out but only with RtcB sequences that belong to cluster 2 of SNN.

To generate SSN shown in Supplementary Fig. 1b, it was necessary to perform the calculation using only 25% of total RtcB sequences, randomly selected. This is because if all RtcB sequences were used, the SSN file would have been too big to be calculated and displayed by Cytoscape using the parameters shown in Supplementary Fig. 1b. Again, SSN file was processed and displayed with Cytoscape, and yFiles Organic Layout was used for the layout of nodes and edges. Minor adjustments of the positions of a couple of clusters were made to make nodes fit better within the space of the figure. The entire SSN was arbitrarily divided into eight clusters, with bacterial RtcB from cluster 2 the main focus of the study.

### Cloning, overexpression, and purification of recombinant proteins

The gene encoding *Ec*PrfH (from *E. coli* ATCC 25922) was cloned into pRSF-1 vector, which carries a N-terminal 6xHis tag followed by a SUMO tag. To obtain *Ec*PrfH with the side chain of Glutamine in the GGQ motif methylated, *Ec*PrfH was co-expressed with methyltransferase PrmC in *E. coli* BL21 (DE3) strain at 18 °C overnight induced with 0.5% lactose^41^. Cells were harvested with centrifugation and the pellets were resuspended in lysis buffer (20 mM Tris-HCl, pH 8.0, 500 mM NaCl, 5% glycerol). Cells were lysed using French Press, and cell debris was removed by centrifugation at 20,000 g for 40 min at 4 °C. The supernatant was filtered with a 0.45 μm filter, and the filtered solution was loaded into a HisTrap column (GE Healthcare). The proteins were eluted with the imidazole gradient. The fractions containing the SUMO-tagged PrfH were combined and incubated with Ulp1 protease to cleave the SUMO tag. The untagged *Ec*PrfH was obtained by passing through the second HisTrap column.

The genes encoding *Ec*RtcB1 and *Ec*RtcB2 were cloned into pETDuet-1 vector without the N-terminal His tag. The two proteins were overexpressed in *E. coli* BL21 (DE3) at 18 °C overnight induced with 0.5 mM IPTG. The harvested cells were resuspended in DEAE-A buffer (20 mM Tris-HCl, pH 8.0, 10 mM NaCl, 5% glycerol) and lysed using French Press. Cell debris was removed by centrifugation at 20,000 g for 40 min at 4 °C, followed by filtration through 0.45 μm filter to clarify the lysate. The proteins were purified from the clarified lysate using an FPLC system with DEAE, Heparin and Superdex 200 size exclusion chromatography. Overexpression and purification of ColD-CT was carried out by following the published procedure^42^.

### Purification of the intact and the damaged *E. coli* 70S ribosome

The intact 70S ribosome was purified using the method described previously^43^. For the damaged 70S ribosome, *E. coli* MG1655 harboring plasmid pCH450::cdiA-CT^ECL^ and pTrc99a::cdiI^ECL^ (both plasmids were generous gifts from C. S. Hayes, UC Santa Barbara) was grown in LB to an OD_600_ of 0.4. 0.2 % arabinose was added to induce the expression of CdiA-CT^ECL^. After 5 min of expression, cells were harvested, and the damaged 70S ribosome was purified analogous to the purification of the intact 70S ribosome.

### Peptide release assays

The kinetic experiments on peptide release were carried out as previously described^41^ with minor modifications. First, the intact 70S ribosomal complexes or the damaged 70S ribosomal complexes, containing intact or cleaved *E. coli* 70S ribosome, f-[^35^S]-Met-tRNA^fMet^, and one of the three mRNAs shown in Fig. 1a, was assembled as previously described. The complex (50 nM) then reacted with *Ec*PrfH of various concentrations at 37 °C. The reactions were quenched by addition of 5% ice-cold trichloroacetic acid at different time points, and the precipitants were removed with centrifugation at 18,000 g for 10 min at 4 °C to separate f-[^35^S]-Met from f-[^35^S]-Met-tRNA^fMet^. The supernatant was recovered and the released f-[^35^S]-Met was counted in 2 mL of Bio-Safe II Complete Counting Cocktail (Research Products International). The maximum releasable fMet (fMet_Max_) was determined by incubating the 70S ribosomal complexes (50 nM) with 200 μM puromycin (Sigma-Aldrich) at 37 °C for 30 seconds^44^. The fraction of f-[^35^S]-Met was determined by the ratio between the released f-[^35^S]-Met from the reaction and fMet_Max_.

### Electron microscopy, data collection and image processing

For the damaged 70S•PrfH, ribosomal complexes were formed by incubating 0.5 μM damaged 70S ribosome with mRNA (1 μM), tRNA^fMet^, and PrfH (5 μM) in buffer A (20 mM HEPES, pH 7.4, 15 mM magnesium acetate, 150 mM potassium acetate, 4 mM β-mercaptoethanol, 2 mM spermidine, 0.05 mM spermine) for 30 min at 37 °C. Sample preparation for cryo-EM was carried out as described^28^. Complexes were first diluted to 80 nM and aliquots of 2.5 μL were incubated for 30 sec on glow-discharged holey carbon grids with thin-layer carbon film (Quantifoil R1.2/1.3, 300 mesh, Copper). Grids were blotted for 3 sec in 100 % humidity at 4 °C and plunge frozen with a Vitrobot (FEI). Data was collected in vitreous ice using a Titan Krios G3i electron microscope operating at 300 keV equipped with a Falcon III direct electron detector (FEI). 4,106 micrographs showing Thon ring beyond 3.5 Å were used. The drifts of movie frames were corrected using MotionCor2^45^, and the contrast transfer functions were determined using CTFFIND4^46^.

Subsequently, a total of 997,751 particles were picked automatically in Relion3 software using Laplacian-of-Gaussian, extracted, and subjected to reference-free 2D classifications to remove non-ribosomal particles^47^. 3D classification was employed to remove the ribosomal subunits. 243,376 particles with PrfH in the A site and tRNA in the P and E site were further analyzed using focused classification with signal subtraction (FCwSS). A total of 207,600 particles were subjected to the final refinement and yielded a global 3D reconstruction with an overall resolution of 2.55 Å.

To determine whether PrfH is able to bind intact ribosome, 0.5 μM intact 70S ribosome, mRNA (1 μM), tRNA^fMet^ and PrfH (5 μM) were mixed in buffer A and incubated at 37 °C for 30 min. 3,750 micrographs were acquired using the parameters the same as the damaged 70S•PrfH complex. Data processing was also carried out in Relion3. No particle with PrfH was found. Instead, most of ribosome were complexed with E-, P-tRNA and mRNA. A total of 87,508 particles were selected for final refinement and yielded a 3.30 Å reconstruction.

Resolutions were reported based on the gold-standard Fourier shell correlation (FSC) of 0.143 criterion^48,49^. Final map was sharpened by applying a negative B-factor estimated using Relion3^47,48^. Local resolution was estimated using ResMap^50^.

### Model building and refinement

Recently reported cryo-EM structure of the intact *E*.*coli* 70S•ArfA•RF2•tRNA complex, excluding ArfA, RF2 and tRNAs, was used as a starting model for structure refinement^25^. The structure of the rRNA-recognition domain of PrfH was built *de novo* according to the density. The peptidyl-hydrolase domain of PrfH was built using domain 3 of RF2 in the 70S•ArfA•RF2•tRNA complex as the starting model^25^. Initial fitting of protein, tRNA and ribosome subunits were performed in Chimera^51^. The structures were refined in Phenix^52^. The final model was validated using MolProbity^53^. Supplementary Table 1 summarizes refinement statistics for the structure. Maps were visualized using Chimera and figures were generated using Chimera and PyMOL.

### Preparation of tRNA and ribosome substrates for RNA repair

Cleaved tRNA^Asp^ was prepared *in vivo* with the expression of ColE5-CT. Thus, the gene encoding ColE5-CT was cloned into Arabinose-inducible pBAD33 vector and transformed into *E. coli* MG1655 cells. When OD_600_ of the *E. coli* cells reach 0.4, expression of ColE5-CT was induced with the addition of 0.2% of L-Arabinose. After 10 min of induction, cells were harvested with centrifugation, and total tRNAs were isolated using acid guanidium thiocyanate-phenol-chloroform extraction. The cleaved tRNAs, which include cleaved tRNA^Asp^, were further purified by denaturing polyacrylamide gel electrophoresis (DPAGE).

Our previous experience indicated that, for the reason we still do not understand, tRNA^Arg^ substrate prepared *in vivo* was not a good substrate. Therefore, cleaved tRNA^Arg^ was prepared *in vitro* using the recombinant ColD-CT. Total tRNAs (70 μM) isolated from *E. coli* MG1655 were incubated with recombinant ColD-CT (0.21 μM) at 37 °C for 10 min in the presence of Cleavage Buffer (25 mM HEPES, pH 7.0, 25 mM NaCl, 5 mM MgCl_2_, 0.05 mg/mL BSA), and the cleaved products were purified by DPAGE.

Preparation of cleaved 70S ribosome for repair assay was similar to the procedure of obtaining the damaged *E. coli* 70S ribosome described previously with minor modification. First, CdiA-CT^ECL^ was expressed for 30 min instead of 5 min. Second, following the expression of CdiA-CT^ECL^, the immunity protein inhibiting CdiA-CT^ECL^, CdiI^ECL^, was also expressed for 3 min with the induction of 1 mM IPTG. The cells were then harvested, and the pellets were resuspended in Ribosome Dissociation Buffer (20 mM HEPES, pH 7.5, 1 mM MgCl_2_, 200 mM NH_4_Cl, 4 mM β-mercaptoethanol) supplemented with 0.5 mM PMSF and 5.5 μg/mL DNase I. The cells were lysed using French Press and the lysate was clarified with centrifugation at 15,000 g for 30 min at 4 °C. This clarified lysate was used directly for the ribosome repair assays.

### Repair of cleaved RNA substrates and Northern blotting analysis

Repair of cleaved tRNA^Asp^ was carried out with either recombinant *Ec*RtcB1 or *Ec*RtcB2 in a reaction mix containing 50 mM Tris-HCl, pH 7.4, 2 mM GTP, 5 mM MnCl_2_, 20 mM MgCl_2_, 6 μM enzyme, and 2 μM of the cleaved tRNA substrate at 37 °C. Repair of the cleaved tRNA^Arg^ was carried out under identical conditions except the concentrations of the enzyme and the tRNA substrate were five times more. The enzyme was preincubated with all components in the reaction mix except the tRNA substrate at 37 °C for 5 min before the addition of tRNA substrate. 8 μL aliquots were taken out at specific time intervals: 2 min, 5 min, 15 min, 30 min and 60 min. The “No enzyme” control was prepared by taking an aliquot of the reaction mix before the addition of the enzyme, followed by addition of the tRNA substrates. The reactions were quenched by addition of 8 μL DPAGE Loading Buffer. The samples were heated at 95 °C for 3 min, followed by 12% DPAGE analysis for 25 min. After the gel analysis, RNAs were transfer to nylon membrane, followed by hybridization with 5’-^32^P-radiolabeled DNA probes targeting the 3’-half of *E. coli* tRNA^Asp^ and tRNA^Arg^, respectively.

For the repair of the damaged 30S subunit, an estimation of ribosomal RNA concentration was made based on the absorbance of the clarified lysate at 260 nm. The reaction mix consisted of 50 mM Tris-HCl, pH 7.4, 2 mM GTP, 5 mM MnCl_2_, 10 μM enzyme and 1 μM of the damaged or the intact 30S subunit. 75 μL aliquots were taken out at the specific time intervals and quenched by addition of TRIzol reagent (ThermoFisher Scientific). This was followed by extraction of total RNA. 3 μg of RNA from each reaction was analyzed by 4% DPAGE gel for 25 min, followed by Northern blotting using a 5’-^32^P-radiolabeled DNA probe complementary to the 3’ end of *E. coli* 16S rRNA^6^.

All Northern blots were exposed to Phosphor screen, imaged using STORM Imager (GE), and the bands quantified using Image Quant software. The intensities of all bands were adjusted with the bands of background. The percentage of repair for tRNA^Asp^ and tRNA^Arg^ was determined by calculating the intensity of the full-length tRNA band as a percentage of the total band intensity (full-length and cleaved) for each sample. Unlike tRNA substrates, intact 16S rRNA is already present in the repair substrate. Therefore, calculation of the percentage of repair of 30S subunit is a bit more complicated. Thus, a total intensity of each lane was obtained with the addition of the intensities of the intact/repaired 16S rRNA and the 49-nt cleavage product. The ratio of this value was calculated for each sample relative to the “No enzyme” control (The sample in the S lane shown in Fig. 4e, f), and the band intensities in each lane were adjusted with this ratio. Next, the band intensity of the intact 16S rRNA band of the “No enzyme” control was subtracted from each of the adjusted repaired 16S rRNA bands. Finally, the extent of repair was determined by calculating the percentage of this repaired 16S rRNA band intensity with respect to the total adjusted band intensities (repaired band and cleavage product).

## Data availability

Electron microscopy map has been deposited in the Electron Microscopy Data Bank under accession codes EMD-24944 for the damaged 70S•PrfH complex. Coordinates have been deposited in the Protein Data Bank under accession codes 7SA4 for the damaged 70S•PrfH complex.

## Acknowledgement

This work was supported by the National Institutes of Health (grant GM120764 to R.H.H.). We thank the cryo-EM center of Southern University of Science and Technology for cryo-EM data collection. We thank C. S. Hayes (UC Santa Barbara) for providing us the plasmids that encodes CdiA-CT^ECL^ and its immunity protein. We also thank P. Wang for carrying out the initial stage of research.

## Author contributions

R.H.H. conceived of the project. Y.T. performed the peptide release assays assisted by F.Z. and A.C. F.Z. carried out cryo-EM experiments assisted by Q.L. A.R. performed RNA repair assays assisted by S.F. A.C. carried out research at the early stage of the project. R.H.H. supervised the research and experimental design. R.H.H. wrote the manuscript with the input from all authors.

