## Supplementary Information for "Sequential rescue and repair of stalled and damaged ribosome by bacterial PrfH and RtcB"

**a**

| Superkingdom | All RtcB | RtcB2 |
|---|---|---|
| Bacteria | 16,044 | 1,437 |
| Archaea | 1,876 | 0 |
| Eukaryota | 1,049 | 98 |
| Viruses | 241 | 1 |

**b**

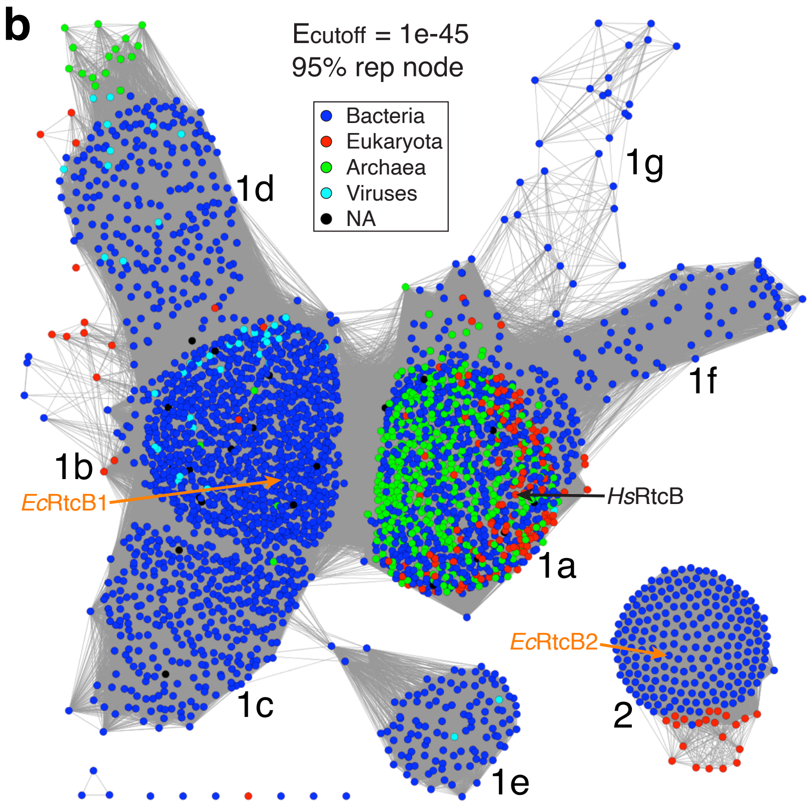


**Supplementary Fig. 1 Bioinformatic analysis of RtcB. a** The numbers of unique sequences of RtcB found in three superkingdoms of life and viruses. **b** Sequence Similarity Network (SSN) of RtcB. See Methods for details of constructing the network. Notably, this SSN was generated using only 25% of the sequences on the database. Each node (colored cycle) represents a collection of RtcB sharing >95% sequence identity (95% rep node). An edge (gray line) connects two nodes if the E-value measuring their sequence similarities is smaller than the cutoff value (1e-45 for this network). The entire SSN has been arbitrarily divided into 8 clusters (1a-g, and 2). The nodes representing two *E. coli* RtcB are marked with orange arrows, and the node representing human RtcB (*Hs*RtcB) is also marked with a black arrow for comparison.

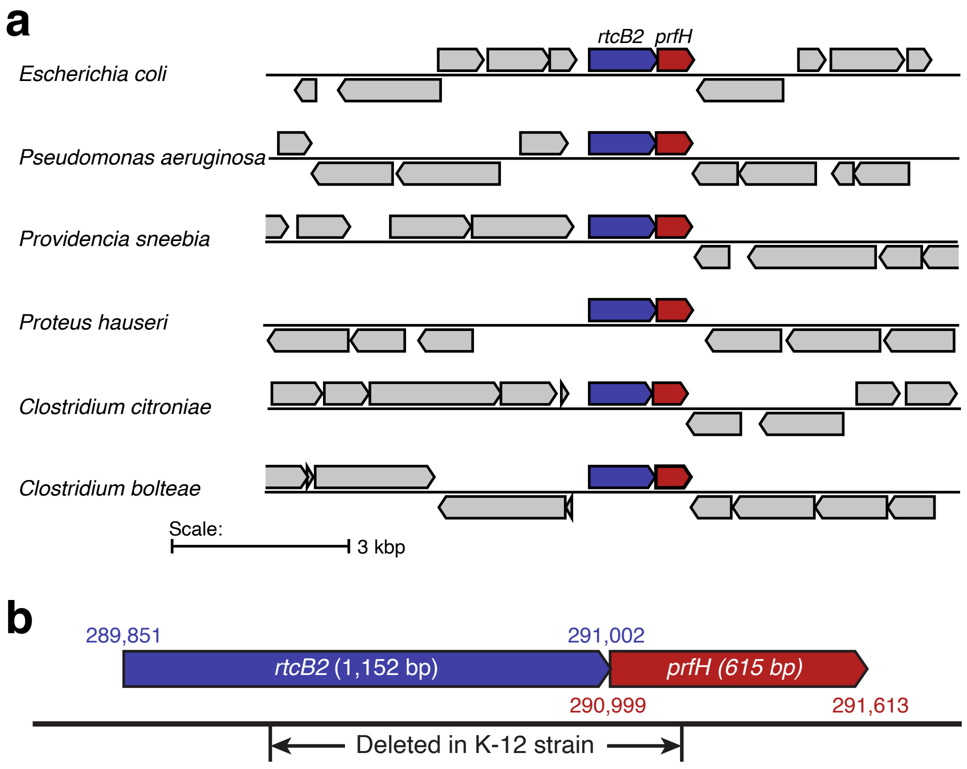


**Supplementary Fig. 2 The genes encoding bacterial RtcB2 and PrfH are predominately found in a two-component operon. a** Six representatives of genomic organizations centered at the *rtcB2-prfH* operon. The genes encoding RtcB2 and PrfH are highlighted in colored boxes. **b** Detailed view of the operon encoding *E. coli* RtcB2 and PrfH. *Ec*RtcB2 and *Ec*PrfH are co-translational, demonstrated by the overlap of the stop codon for RtcB2 and the start codon for PrfH. All *E. coli* strains encode full-length RtcB2 and PrfH except for K-12 strain and its derivatives, which have a large genomic deletion that encompasses the coding regions of both proteins.

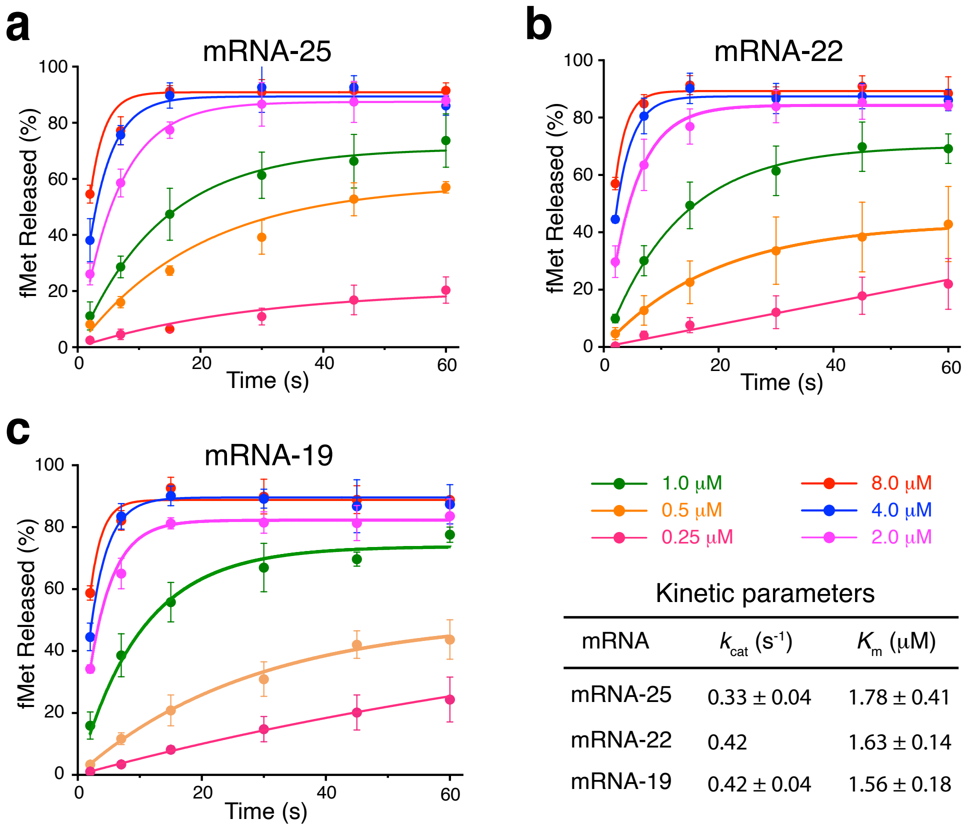


Kinetic parameters

| mRNA | $k_{cat}$ ($s^{-1}$) | $K_m$ (μM) |
|---|---|---|
| mRNA-25 | 0.33 ± 0.04 | 1.78 ± 0.41 |
| mRNA-22 | 0.42 | 1.63 ± 0.14 |
| mRNA-19 | 0.42 ± 0.04 | 1.56 ± 0.18 |

**Supplementary Fig. 3 Expanded peptide release assays with *Ec*PrfH.** (**a-c**) Time course of the peptide release assays using various concentrations of *Ec*PrfH. Each experiment was repeated three times, and the same assays were carried out for three different mRNAs listed in Fig. 1a. These assays allowed us to obtain kinetic parameters listed in the table.

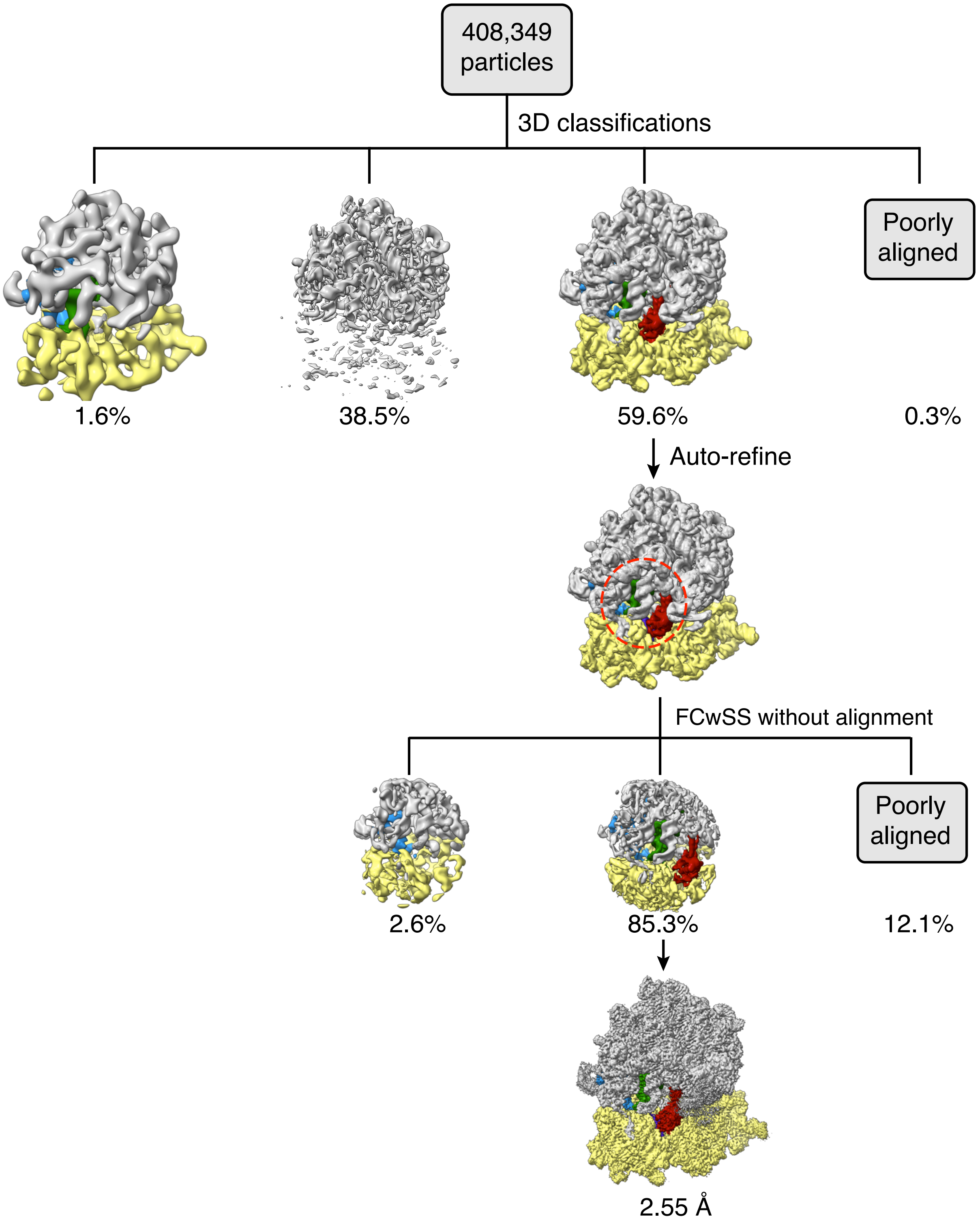


**Supplementary Fig. 4 Cryo-EM classification of *Ec*PrfH in complex with the damaged *E. coli* 70S ribosome.** The complete dataset of 408,349 particles was initially classified in four classes. The resulting class 2 (243,376) was further refined and subject to focused classification with signal subtraction (FCwSS) in Relion. The remaining 207,600 particles containing PrfH were then 3D-refined, resulting in a final reconstruction of 2.55 Å average resolution.

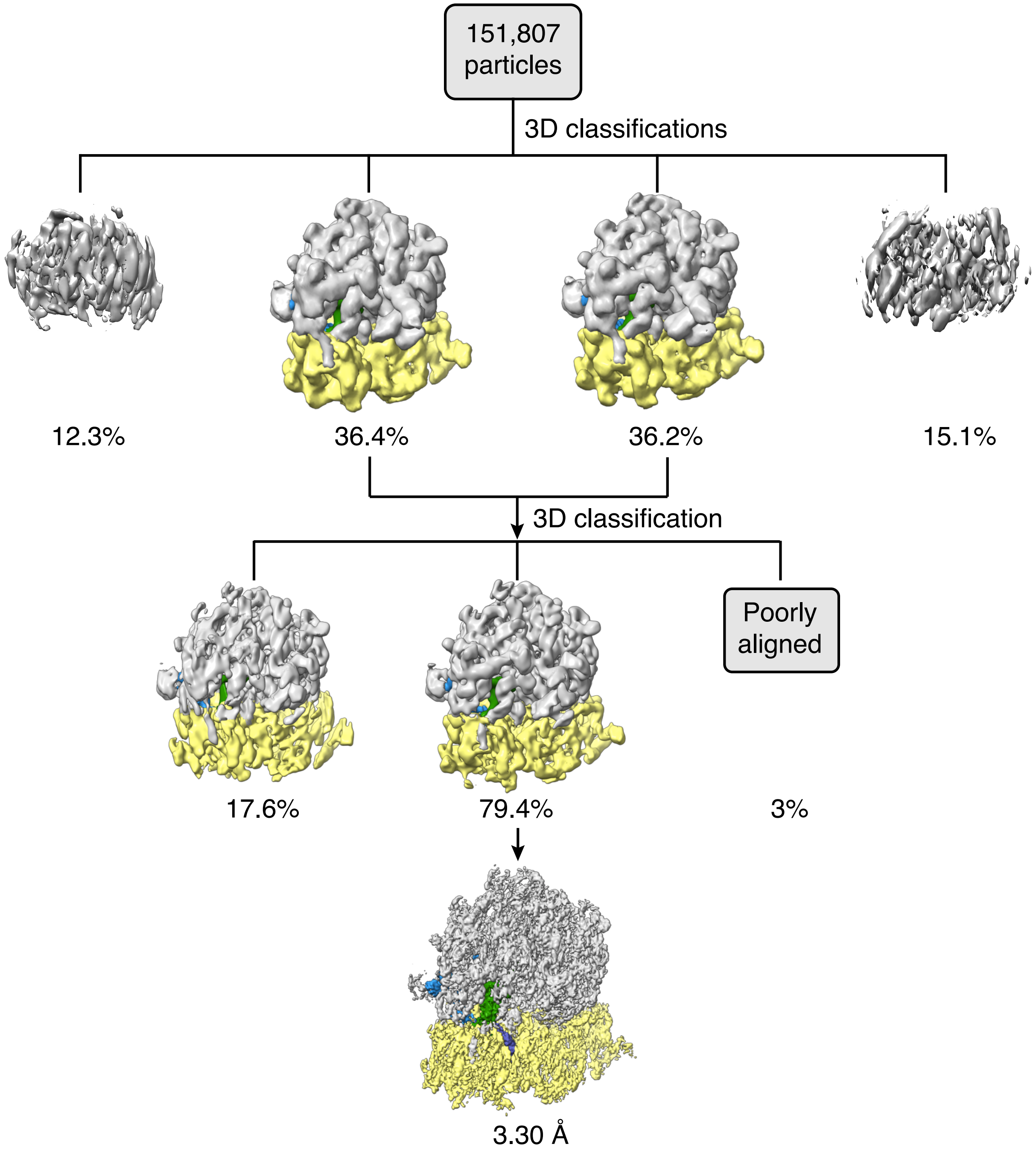


**Supplementary Fig. 5 Cryo-EM classification of *Ec*PrfH in complex with the intact *E. coli* 70S ribosome.** 151,807 particles selected from 2D classification were classified by two rounds of 3D classifications in Relion to removing the subunits and poorly aligning particles. The final reconstruction with P- and E-tRNA in the ribosome has a nominal resolution of 3.3 Å.

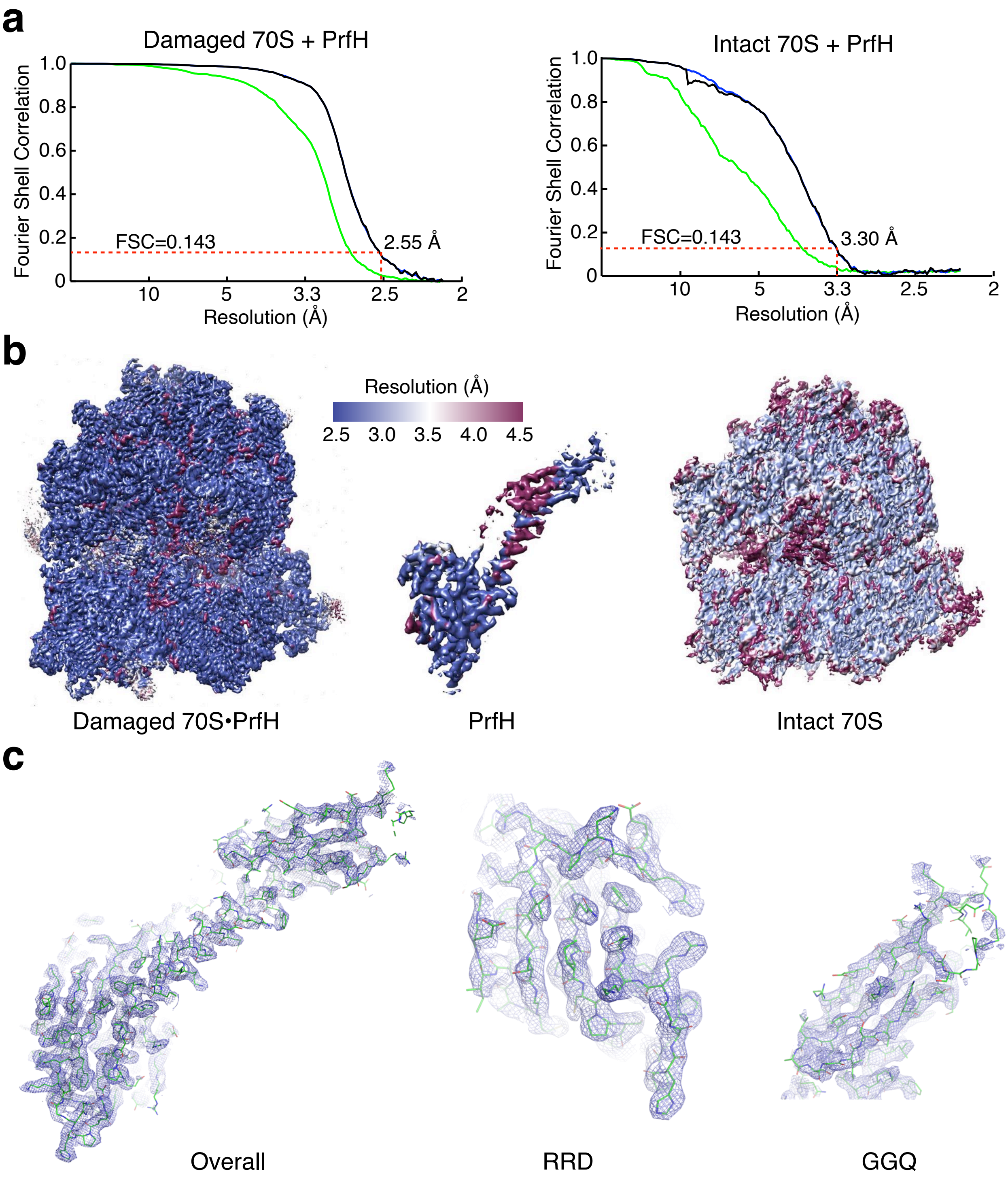


**Supplementary Fig. 6 Cryo-EM resolution and map.** **a** Gold-standard FSC curves for the electron microscopy map of damaged 70S ribosome (left panel) and intact 70S ribosome (right panel). Resolution is demarcated using the FSC=0.143 criterion. **b** Unfiltered and unsharpened density map colored by local resolution in surface for the damaged 70S ribosome, PrfH, and the intact 70S ribosome. **c** Representative electron microscopy maps showing the refined structures of the overall PrfH, the rRNA-recognition domain, and the peptide-hydrolase domain.

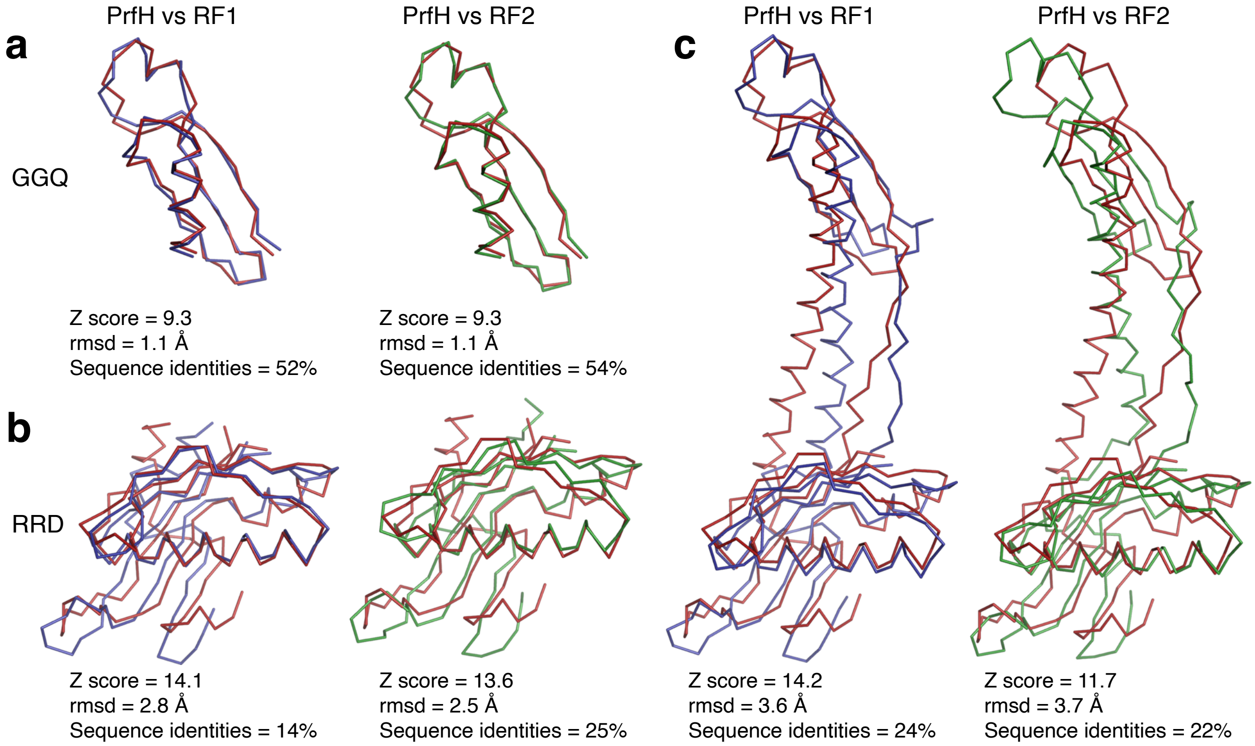


**Supplementary Fig. 7 Structural alignments of the individual domains (a, b) as well as the full-length protein (c) of *Ec*PrfH with their counterparts in RF1 and RF2.** The pairwise structural comparisons were carried out at Dali server (http://ekhidna2.biocenter.helsinki.fi/dali/). The Z score of Dali search as well as rmsd of the three structures are listed. The pairwise sequence alignments were carried out using Blastp at NCBI, and the percentiles of sequence identities are also listed. PrfH is colored red, RF1 is in blue, and RF2 is in green.

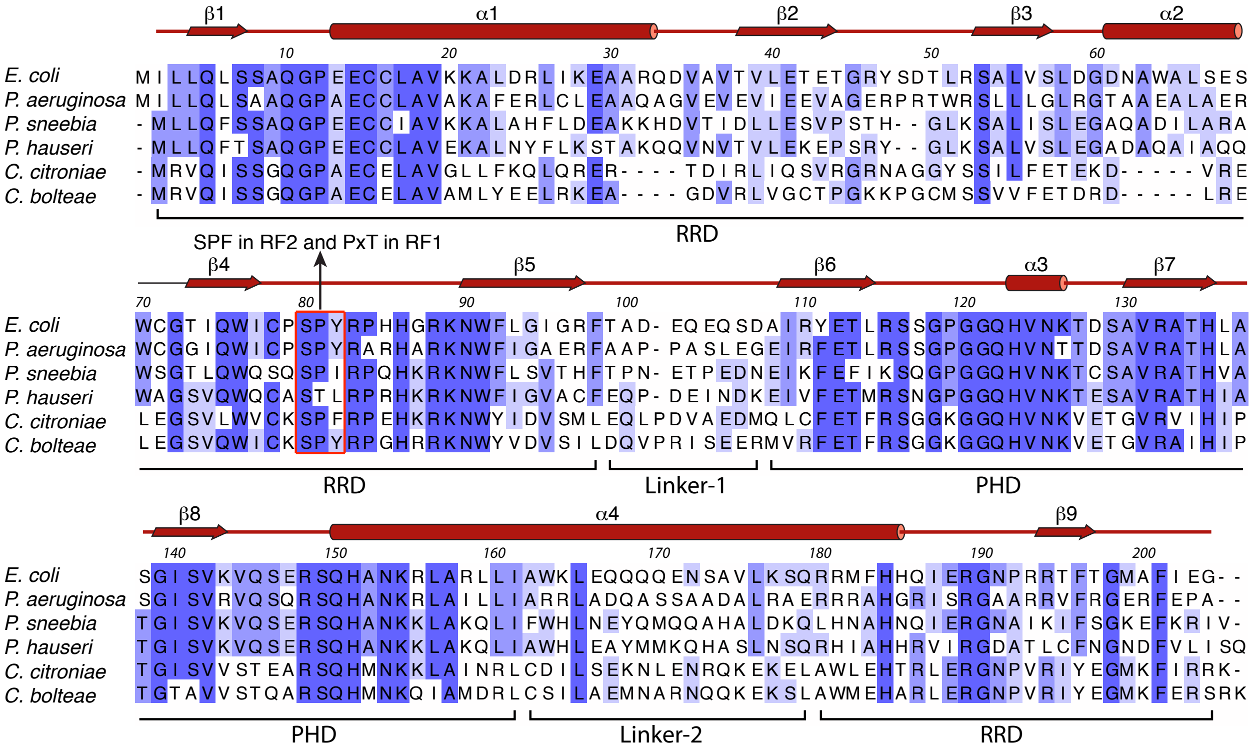

**Supplementary Fig. 8 Secondary structure and domain boundaries of PrfH.** PrfH sequences from six organisms shown in Supplementary Fig. 2a were aligned. The conservation was colored by different shades of blue. The numbers on the top of the sequences correspond to *Ec*PrfH. The secondary structures were depicted at the top, and the domain boundaries were marked at the bottom. The triple residues corresponding to the most conserved motifs in RF1 and RF2 are boxed in red.

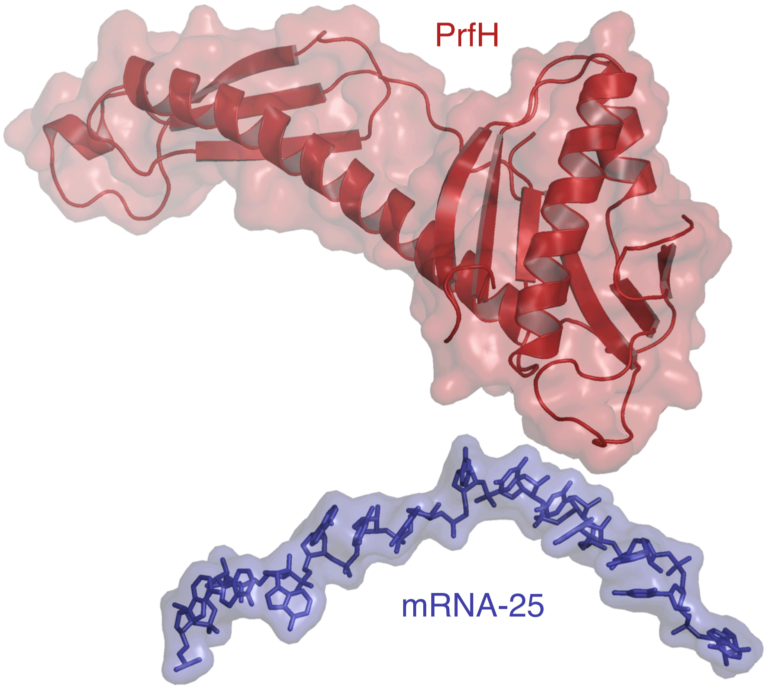


**Supplementary Fig. 9 PrfH and mRNA do not make contacts.** Structure of the damaged *E. coli* 70S•PrfH•tRNA•mRNA complex with only the structures of PrfH and mRNA showing for clarity. PrfH is depicted in cartoon and surface, and mRNA is depicted in sticks and surface.

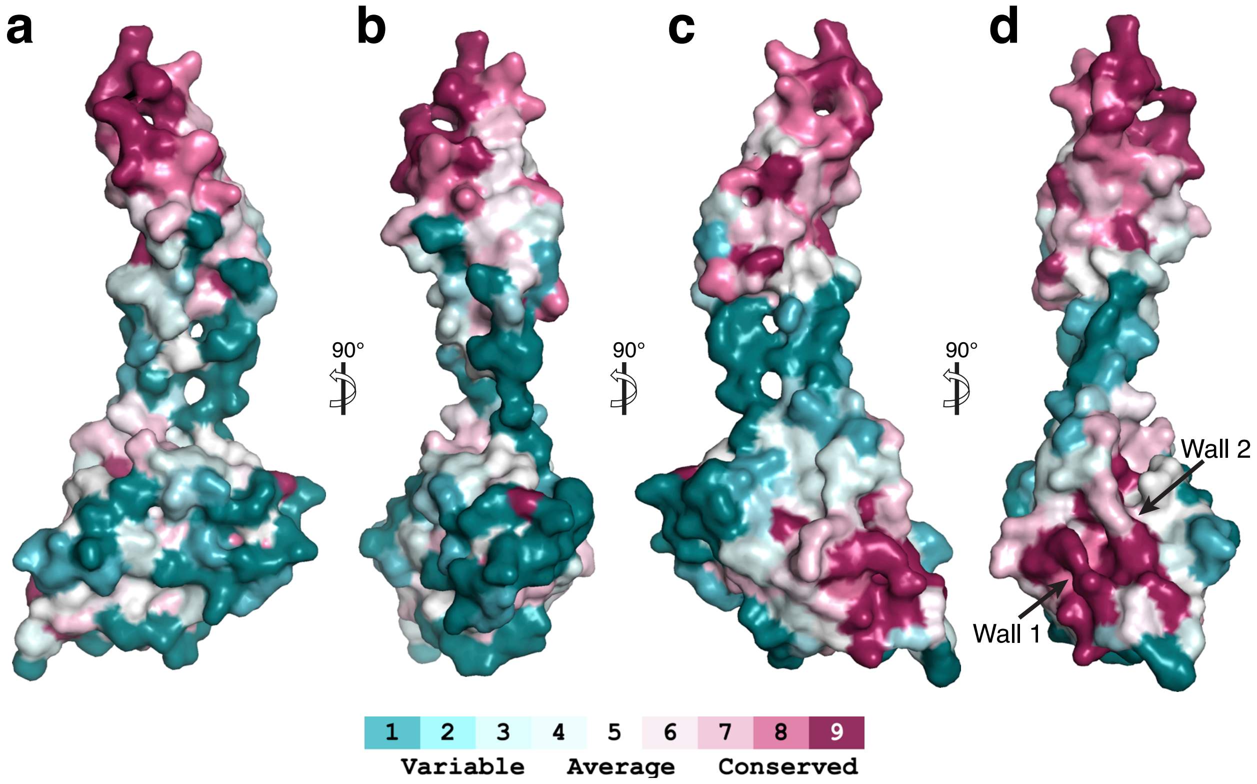


**Supplementary Fig. 10 Mapping the conservation of amino acid sequences of PrfH on the structure of *Ec*PrfH.** The sequence of *Ec*PrfH and additional 200 randomly selected PrfH sequences were aligned. The aligned PrfH sequences, together with the structure of *Ec*PrfH, were employed for ConSurf analysis, resulting in the conservation of each residue to be assigned a number (1-9). The *Ec*PrfH structure is depicted in surface, with each residue colored by the degree of conservation. The structures in panels **a** and **b** have the same orientation as the ones in Fig. 2b. The peptide-hydrolase domain (top) has the highest conservation, followed by the rRNA-recognition domain (bottom). The linker regions are least conserved. The data shown here are consistent with the sequence alignments shown in Supplementary Fig. 8.

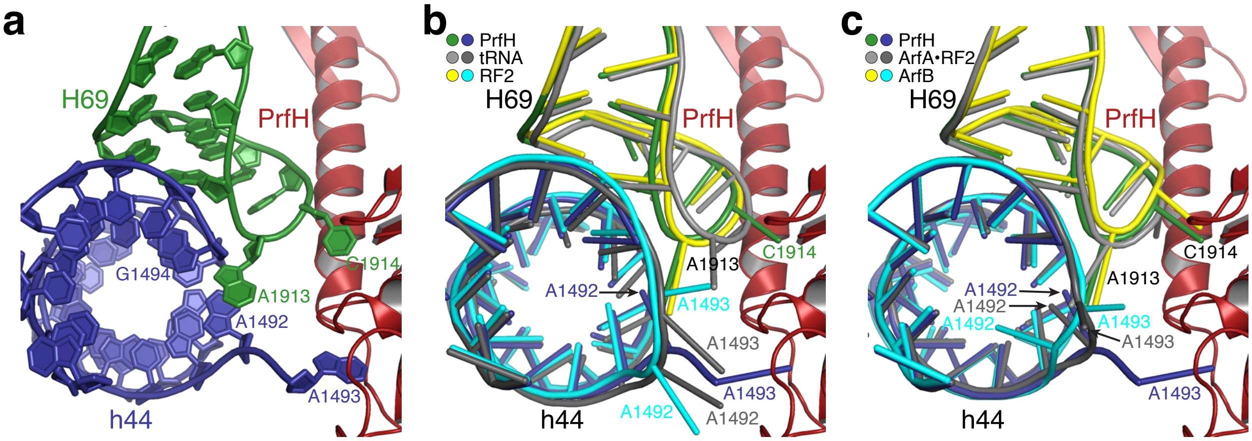


**Supplementary Fig. 11 Conformational changes of ribosomal RNAs resulting from PrfH bound to the damaged 70S ribosome. a** Cartoon representation of the structure viewing into the helix of h44. The ribose and the base of the nucleotides were depicted in plates. Both A1493 and C1914 are flipped out from helices and flipped into their respective recognition pockets in PrfH. A1913 occupies the space vacated by extrahelical A1493, stacking on top of A1492. **b** Cartoon representation of the structures of 70S ribosome with PrfH, RF2, and tRNA occupying the A site. Nucleotides are depicted with rods to facilitate comparisons. **c** Cartoon representation of the structures of 70S ribosome with PrfH, ArfB, and ArfA•RF2 occupying the A site. Similarities among these three structures include: (i) All A1492 are interhelical; (ii) All A1493 are extrahelical; (iii) All A1913 are in similar locations; and (iv) All C1914 are extrahelical.

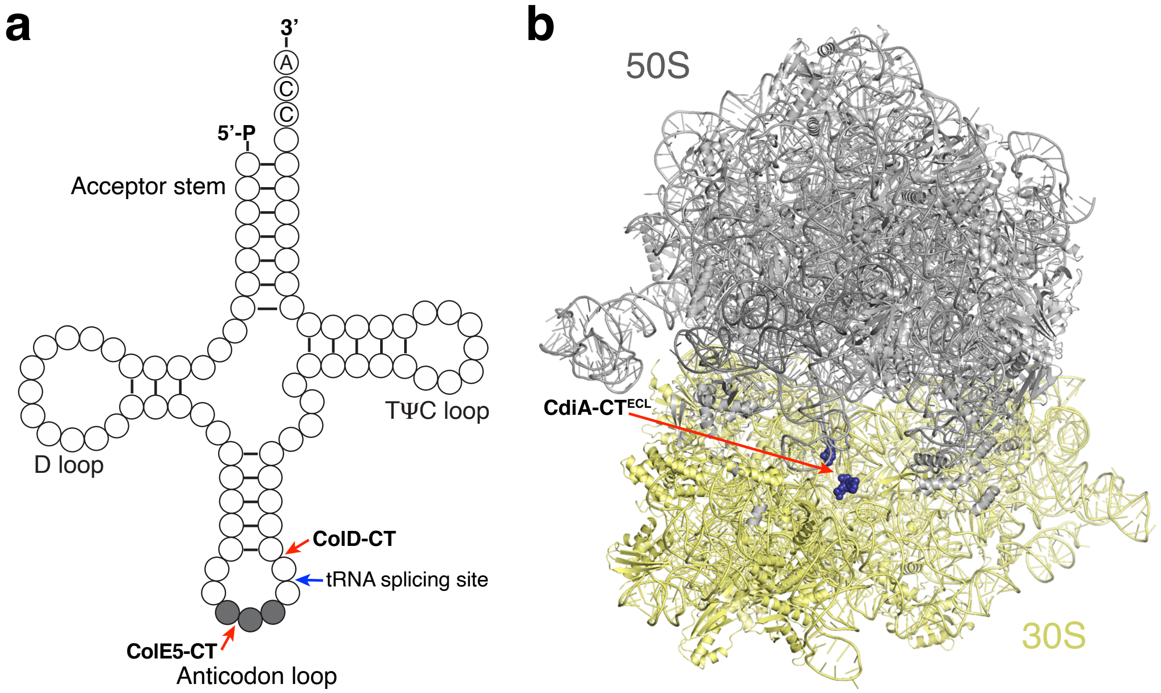


**Supplementary Fig. 12 Schematic view of three RNA substrates employed for RNA repair by *Ec*RtcB1 and *Ec*RtcB2. a** Cloverleaf view of tRNA. The anticodons are colored gray. The sites of cleavage by ColE5-CT and ColD-CT are marked with red arrows. The tRNA splicing site, which occurs in archaeal and eukaryotic organisms, is marked with a blue arrow for comparison. **b** the cryo-EM structure of the damaged *E. coli* 70S ribosome in complex with PrfH (not shown). The structure is depicted the same as in Fig. 1a. The two terminal nucleotides at the cleavage site, A1493P and G1494, are depicted in sphere and colored blue. The conformation of A1493P might be different in the free form of the damaged 70S ribosome.

**Supplementary Table 1 Cryo-EM data collection and model statistics**

| | Damaged 70S•PrfH complex |
|---|---|
| **Data Collection** | |
| Particles | 997,751 |
| Pixel size (Å) | 1.05 |
| Defocus range (μm) | -0.6 to -3.0 |
| Voltage (kV) | 300 |
| Electron dose ($e^{-}$Å$^{-2}$) | 30 |
| **Model composition** | |
| Non-hydrogen atoms | 150,065 |
| Protein residues | 6,175 |
| RNA bases | 4,724 |
| Ligands ($Zn^{2+}$/$Mg^{2+}$) | 2/439 |
| **Refinement** | |
| Resolution (Å) | 2.6 |
| $FSC_{average}$ | 0.90 |
| **Rms deviations** | |
| Bond lengths (Å) | 0.010 |
| Bond angles (°) | 0.843 |
| **Validation (proteins)** | |
| MolProbity score | 1.81 |
| Clashscore, all atoms | 11.31 |
| Favored rotamers (%) | 99.37 |
| **Ramachandran plot** | |
| Favored (%) | 96.34 |
| Outliers (%) | 0.05 |
| **Validation (RNA)** | |
| Correct sugar puckers (%) | 99.51 |
| Good backbone conformations (%) | 83.81 |
